# Multi-parametric spectral modeling disentangles synaptic kinetics and neural excitability from non-invasive electrophysiological recordings

**DOI:** 10.64898/2026.10.15.759313

**Authors:** Vincent Philippe, Lise Tombroff, Fahimeh Akbarian, Sara De Witte, Nicola Trotta, Jeroen Van Schependom, Sebastiaan Engelborghs, Jean-Christophe Bier, Mélanie Strauss, Xavier De Tiège, Vincent Wens

## Abstract

Spectral modeling of electrophysiological recordings enables non-invasive characterization of neural excitability. However, current approaches rely on a single spectral index, limiting insight into the multiple biophysical processes underlying excitation-inhibition balance. Here, we introduce a biologically informed, multi-parametric spectral analysis that disentangles neural excitability and synaptic kinetics from recordings. As a proof of concept, we demonstrate that this approach captures signatures of hyperexcitability and glutamate reuptake dysfunction characteristic of Alzheimer’s disease.

## Main text

*In vivo*, neural circuits operate in a regime of balanced excitation and inhibition (E/I), with elevated neural excitability supporting rapid responsiveness.^1^ State-of-the-art approaches to inferring this balance from human electrophysiological recordings, such as magnetoencephalography (MEG), rely on spectral models that separate periodic and aperiodic components^2^ and quantify neural excitability using the exponent of a power-law fit to the aperiodic spectrum.^3^ These methods successfully characterized E/I imbalance in brain disorders including epilepsy,^4^ multiple sclerosis,^5^ autism spectrum disorder,^6^ or Alzheimer’s disease (AD).^7^ However, E/I balance emerges from multiple biophysical processes that cannot be captured by a single spectral parameter. Here, we present a biologically informed, multi-parametric spectral analysis that disambiguates distinct underlying mechanisms.

To describe and validate our approach, we used AD as an experimental model of E/I imbalance, linking synaptic dysfunctions at the microscale circuit level^8^ with accompanying macroscale alterations in glucose metabolism.^9^ We analyzed a dataset of MEG and positron emission tomography with [^18^F]-fluorodeoxyglucose (FDG-PET) in 43 biomarker-positive patients with AD. We first sought to replicate the reported brain-wide association between hyperexcitability and hypometabolism in AD, using the classical neural excitability index defined as the exponent of an aperiodic power-law fit to regional MEG spectra in the 15–50 Hz frequency range (Fig. 1a, blue).^7^ We recovered a significant negative association between this index and regional cerebral metabolic uptake (Fig. 1b; *β* = −0.29 ± 0.08, mean ± SEM). Because the 15–50 Hz range overlaps synaptic timescales, this E/I imbalance likely reflects synaptic dysfunction, with neurocomputational simulations suggesting hyperexcitability arising from an excess ratio of glutamatergic to GABAergic postsynaptic conductances.^3^ However, this interpretation remains constrained by the single-parameter nature of this analysis. We introduce below a multi-parametric extension that refines this mechanistic picture.

**Figure 1:**
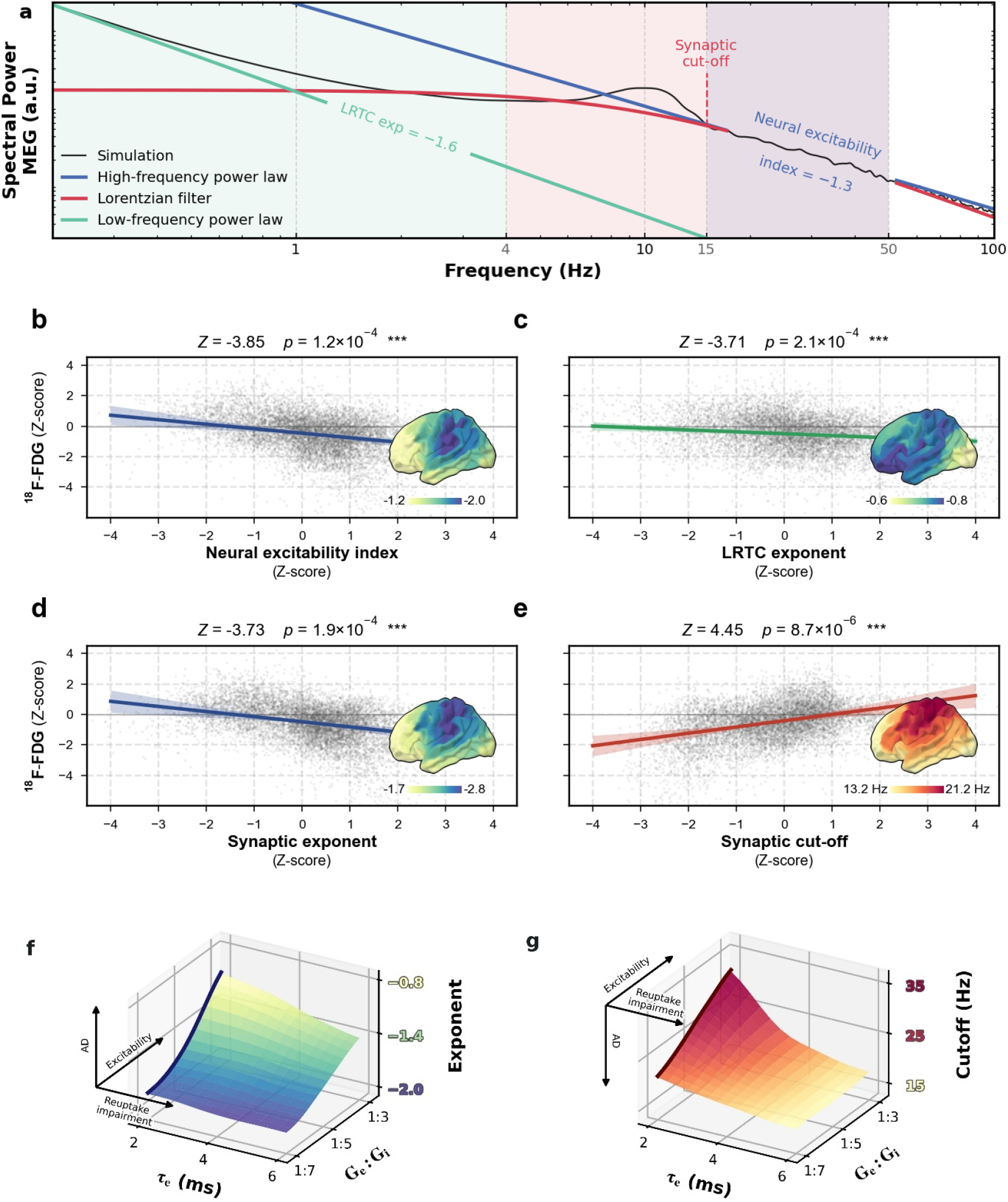
MEG/FDG-PET analysis of AD patients. **a**, Illustration of spectral fits on simulated MEG data including the low-frequency power law measuring network LRTC (green), the Lorentzian measuring synaptic lowpass filtering (red), and the high-frequency power law underlying the neural excitability index (blue). Light-colored backgrounds represent fitting windows. Care was taken to avoid biases due to the *α* oscillation.^2^ **b–e**, Association between^18^F-FDG (standardized uptake value ratio) and each spectral parameter of interest: neural excitability index (b), LRTC exponent (c), synaptic exponent (d) and cut-off frequency (e). All data are from patients with AD after standardization over a normative dataset of twenty-five healthy controls. Inserts show brain maps of each parameter without standardization. Z-scores and *p*-values indicate the result of linear mixed models. **f**,**g**, Simulations assessing synaptic exponent (f) and cut-off frequency (g) parameters as functions of the excitatory-to-inhibitory conductance ratio (*G*_*e*_ : *G*_*i*_) and excitatory time constant (*τ*_*e*_) with fixed inhibitory time constant *τ*_*i*_ = 10 ms, following Gao et al.^3^ Thick curves correspond to the case of AMPA time constant fixed to a baseline physiological value *τ*_*e*_ = 2 ms.^3^ Axis inserts guide the interpretation of parameters.

Balanced E/I yields asynchronous, irregular neuronal firing^10^ that is transduced by synapses into large aperiodic fluctuations in postsynaptic conductances.^1^ These fluctuations span two distinct spectral domains,^11^ motivating a multi-parametric approach. First, the low-frequency domain is dominated by a power law emerging from large-scale, network-level recurrent excitation.^12^ We fit a power-law below 4 Hz to estimate its scaling exponent (Fig. 1a, green), which quantifies long-range temporal autocorrelation (LRTC) in presynaptic firing rates. This electrophysiological low-frequency exponent is particularly interesting because previous work did not probe this domain.^7^ We found a significant association with regional cerebral metabolism in our patients (Fig. 1c; *β* = −0.12 ± 0.03), showing that AD hypometabolism is accompanied with LRTC over shorter timescales. This suggests that AD pathophysiology accentuates the temporal irregularity of network-level neural firing characteristic of E/I balance, consistent with previous fMRI data.^13^

This leads us to the higher-frequency domain of E/I balance (approximately 10–80 Hz), which reflects local transduction of presynaptic spikes into postsynaptic conductances.^11^ Within this domain, synaptic transduction acts as a combination of first-order lowpass filters controlled by the relative conductances and kinetics of AMPA and GABA_A_ receptors.^3,11^ While MEG studies typically rely on high-frequency power-law characterizations (as with the neural excitability index),^2,7^ we adopted a more principled strategy drawn from intracranial recordings,^14^ fitting a Lorentzian law to our regional MEG spectra in the 4–50 Hz range (Fig. 1a, red). Lorentzian curves provide minimal models of synaptic lowpass filtering, parameterized by a cut-off frequency reflecting the presynaptic spike rate above which transduction loses efficiency and by a scaling exponent reflecting the sharpness of attenuation. Both parameters were associated with cerebral metabolic uptake; specifically, AD hypometabolism related to higher exponent and lower cut-off frequency (Figs. 1d,e; *β*_exponent_ = −0.33±0.09, *β*_cut-off_ = 0.41±0.09). Our two-parameter Lorentzian fit provided additional information not reducible to the neural excitability index alone, significantly improving the association with regional cerebral hypometabolism (nested model comparison, difference in Akaike information criterion ∆AIC = −4.4, *χ*^2^(2) = 8.43, *p* = 0.015). Moreover, the neural excitability index was largely explained as an aggregate of the two synaptic parameters (*β*_exponent_ = 1.13, *β*_cut-off_ = 0.41; *R*^2^ = 0.75, *p <* 10^*−*10^). The synaptic exponent and cut-off frequency therefore capture part of the regional hypometabolism that the neural excitability index alone misses, providing a closer link to underlying pathophysiological mechanisms of E/I imbalance.

To gain mechanistic insight into what the synaptic exponent and cut-off frequency represent in terms of circuit-level biophysics, we simulated ensembles of excitatory AMPA and inhibitory GABA_A_ synapses in E/I balance^3^ and computed our two spectral parameters across varying E/I conductance ratios. Hyperexcitability increased both exponent and cut-off (Figs. 1f,g; thick curve), revealing an apparent paradox. While the conventional picture of hypometabolism/hyperexcitability in AD matched the exponent (both increased the exponent, like the neural excitability index), it was inconsistent for the cut-off frequency on which regional cerebral hypometabolism in AD and simulated hyperexcitability had opposite effects. Resolving this discrepancy required modulating the AMPA receptor time constant in our simulations, which revealed a clear mapping: the synaptic exponent primarily tracks the E/I conductance ratio (Fig. 1f), whereas the cut-off frequency reflects transduction kinetics, with lower cut-offs indicating longer excitatory time constants (Fig. 1g). Together, this suggests a refined picture in which regional cerebral hypometabolism in AD is consistent with both hyperexcitability and prolonged postsynaptic glutamatergic signaling. This echoes impaired glutamate reuptake observed in transgenic mouse models of AD,^8^ leading to longer glutamate dwell times in excitatory synapses potentially captured by our cut-off frequency parameter. In conclusion, multi-parametric spectral modeling of MEG recordings goes beyond neural excitability, providing access to synaptic mechanisms of E/I balance that are normally confined to invasive, circuit-level approaches.^14^ Our method includes three interpretable parameters: the synaptic exponent, tracking intrinsic neural excitability; the cut-off frequency, tracking synaptic kinetics; and the LRTC exponent, tracking emergent network autocorrelations. This is supported by the association of the synaptic parameters and of the LRTC exponent with regional cerebral hypometabolism in AD, the synaptic parameters adding information to the neural excitability index, and by the consistency with known cellular mechanisms of AD synaptic dysfunction in mouse models. By enabling inference of microscopic biophysical parameters from non-invasive extracranial electrophysiology, our multi-parametric spectral approach offers a transformative opportunity to link macroscale recordings to microscale circuit mechanisms directly *in vivo*.

## Methods

### Dataset description

Participants. Forty-three patients (age, 72 ± 7 years, mean ± SD; 29 females) were recruited from the neurology departments of HUB–Hôpital Erasme (*n* = 25) and UZ Brussels (*n* = 18) for a multimodal study on AD including structural magnetic resonance imaging (MRI), FDG-PET, and resting-state MEG. All patients were biomarker positive (cerebrospinal fluid analysis, *n* = 37; amyloid PET, *n* = 6) and ranged clinically from the preclinical stage to early dementia due to AD according to the Clinical Dementia Rating score (0.8 ± 0.4; normal cognition, *n* = 2; mild cognitive impairment, *n* = 28; early dementia, *n* = 13) and the Mini Mental State Examination score (24.0 ± 3.8). We also used a normative group of 25 healthy controls (age, 65 ± 8 years; 18 females) taken from a previously published dataset.^15^ Our cohort of patients with AD exhibited typical cerebral glucose hypometabolism at the posterior temporoparietal junction and the precuneus (Fig. S1) as well as hippocampal and hippocampal-amygdalo-ventricular atrophy (*t* tests on percentage deviation from normative volume^16^ with age as covariate of non-interest, *p <* 10^*−*6^). Experimental protocols were approved beforehand by local ethics committees of both recruiting institutions (HUB–Hôpital Erasme, P2017/427, EudraCT/CCB: B4062022000111; UZ Brussel, BUN 1432023000065). All participants gave written informed consent prior to their inclusion.

MRI and FDG-PET. Both structural MRI and FDG-PET scans were acquired at HUB–Hôpital Erasme on a hybrid PET-MR system (3T Signa, General Electric). Structural 3D T1-weighted MRIs were used for spatial normalizations, brain parcellation based on the Brainnetome atlas^17^ restricted to cortical regions, and MEG head modeling. Individual FDG-PET images were reconstructed using ordered subset expectation maximization with weighted attenuation correction and 4-mm smoothing. Anatomical normalization was based on a PET template in MNI space built from the target patients’ FDG-PET images.^18^ We used the standardized uptake value ratio referenced to individual pons for intensity normalization and Brainnetome atlas parcels for regional quantification of cerebral glucose metabolism.

MEG. Wakeful resting-state MEG was recorded for 5 minutes with a 306-channel whole-head neuromagnetometer (Triux, MEGIN; acquisition band-pass, 0.1–330 Hz; sampling rate, 1 kHz) installed at HUB–Hôpital Erasme. Subjects were seated with eyes open to minimize fluctuations in wakefulness and in the posterior *α* rhythm that possibly confound spectral analyses. Environmental noise and movement artefacts (tracked by head position indicator coils) were suppressed using spatiotemporal signal-space separation (Maxfilter v2.1 with default parameters, MEGIN).^19^ One minute of artifact-free signal (i.e., minimal excessive scatter at magnetometer amplitude *<* 10 pT) was selected, band-pass filtered (0.1–50 Hz), subjected to independent component analysis to remove cardiac and ocular artefacts,^20^ and source projected by beamforming^21^ (input covariance computed from preprocessed resting-state signals; source space, tangential dipole basis on 5-mm grid defined in MNI space and normalized onto individual MRI;^22^ forward model based on MEG-MRI co-registration and single-layer boundary element head model^23^ derived from MRI tissue segmentation^24^). Regional power spectral density (PSD) in each cortical parcel of the Brainnetome atlas was computed for the reconstructed source with maximum variance within each parcel, using Welch’s method (5-s windows, 50% overlap).

### Spectral analyses

Neural excitability index. The neural excitability index was defined as Ranasinghe et al. as the scaling exponent of a power law fitted to regional PSDs over the 15–50 Hz frequency range^7^ using the “fitting-oscillations-and-one-over-f” (FOOOF) algorithm (FOOOF toolbox v1.1), which excludes frequency bins in the presence of oscillatory peaks in the fitting range.^2^

We describe below the multi-parametric generalization developed here and based on a Lorentzian fit to regional PSDs in synaptic timescales followed by a power-law fit to residuals at low frequencies. We emphasize the original elements that were key to our analysis.

High-frequency Lorentzian fit (synaptic filtering). We implemented a Lorentzian fit to the PSD as a function of frequency *f* according to the parameterization

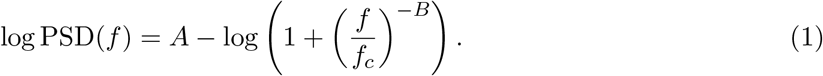

The cut-off frequency parameter *f*_*c*_ corresponds to the characteristic frequency at which the Lorentzian PSD is halved. The exponent *B* controls the approximate power-law decay at high frequencies (*f* ≫ *f*_*c*_). *B* is the log-log slope of the decay, larger values mean a flatter decay. Replacing the high-frequency power law of the neural excitability index with a Lorentzian yields a more principled description of synaptic lowpass filtering and, crucially, provides access to the cut-off frequency.^14^ To limit the effect of oscillations, the Lorentzian was fitted in two steps, as for the neural excitability index: a first least-squares fit on the log PSD, removal of the frequency bins that contained an oscillatory peak, and a second fit without those bins.

A key technical aspect concerned the selection of the 4–50 Hz frequency range for this Lorentzian fit (instead of 15–50 Hz used by Ranasinghe et al.). The 4 Hz lower bound resulted from a compromise between two opposing constraints. Appropriate Lorentzian fit requires probing the PSD reasonably below the expected range of cut-off *f*_*c*_ values. Typical time constants *τ* of fast synapses reach up to 20 ms,^25^ corresponding to 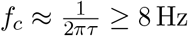 Hz, so the fit domain should extend below 8 Hz. However, it could not be extended too far lest the power law that dominates at low frequencies (*<* 1 Hz) biases the Lorentzian fit, pushing *f*_*c*_ to 0 and reducing the Lorentzian to a power law. The 50 Hz power line bins were removed beforehand. Neurocomputational simulations described below provide further check that expected cut-off frequencies *f*_*c*_ remain well below 50 Hz. The appropriateness of our fit domain was also empirically supported by an examination of residual spectra (Fig. S2).

Low-frequency power-law fit (LRTC). To assess long-range temporal correlations (LRTC) in MEG signals, we extracted the residuals of the Lorentzian fit and fitted a low-frequency power law in the 0.2–4 Hz domain using logarithmic regression. The LRTC exponent was defined as the signed log-log slope of this power law, so that a larger exponent means a flatter spectral decay. Statistics. To examine associations between MEG spectral indices and FDG-PET uptake across brain regions, all regional data in patients with AD were first *z* scored according to the corresponding distribution in control subjects (allowing to suppress inter-regionality variability and to use the 210 cortical Brainnetome regions as repeated measures), and then submitted to hierarchical linear mixed-effect models (LMMs; dependent variable, FDG *z* score; independent variable, spectral index *z* score; random intercept and random slope per patient; age and parcel as fixed-effect covariates).^7^ The added value of the Lorentzian parameters was tested by comparing two nested LMMs fitted by maximum likelihood: the neural excitability index alone, versus the index plus the Lorentzian parameters: *f*_*c*_ and *B*. Both models had the same random effects (random intercept and random slopes for the four indices per patient), so that AIC differences only reflect the fixed effects, and were compared with a likelihood-ratio test (*χ*^2^; degrees of freedom, number of added parameters). We also evaluated whether the neural excitability index could be explained as a combination of the Lorentzian parameters *f*_*c*_ and *B*. We used an ordinary least-squares regression across all regions, with the neural excitability index *z* score as dependent variable and the cut-off *f*_*c*_ and exponent *B z* scores as independent variables and report its *R*^2^ and the *p*-value of the corresponding *F*-test.

### Neurocomputational simulations in E/I balance

Neurocomputational simulations under E/I balance followed the framework of Gao et al., in which Poisson trains of presynaptic spikes were converted into a MEG-like post-synaptic current by convolution with difference-of-exponential functions (parameterized by a fast rise and a slower decay time constant) modeling excitatory (AMPA) and inhibitory (GABA_A_) synaptic transduction, multiplication by a conductance gain parameter (AMPA, *G*_*e*_; GABA_A_, *G*_*i*_) and by the synaptic channel driving force, and summation over excitatory and inhibitory currents.^3^ We set the decay time constants to *τ*_*e*_ = 2 ms for AMPA synaptic excitation and *τ*_*i*_ = 10 ms for GABA_A_ fast inhibition, following Gao et al. framework,^3^ in which synaptic time constants are drawn from CNRGlab @ UWaterloo.^25^ The E/I ratios *G*_*e*_ : *G*_*i*_ were varied from 1:7 (low excitability) to 1:3 (high excitability). In our extended simulation, we further let *τ*_*e*_ vary from 2 to 6 ms, effectively modeling impaired glutamate reuptake in AD. For each simulation, we extracted the synaptic cut-off frequency *f*_*c*_ and exponent *B* using our Lorentzian fit within 4–50 Hz. Of note, these simulations led to cut-off frequencies consistently within 10–35 Hz and thus well inside our selected fit domain (4–50 Hz), thereby providing further theoretical support to our empirical validation.

## Data availability

Open access to post-processed regional MEG spectra and regional FDG-PET data analyzed in this study are available at https://osf.io/t753c. Access to raw data will be made available upon request to the corresponding author and following approval of institutional authorities (HUB–Hôpital Erasme, ULB, UZ Brussel, and VUB).

## Code availability

The code for our multi-parametric analysis of regional MEG and FDG-PET data is available on GitHub at https://github.com/ln2t/lorentzian-ad.

## Acknowledgements

V.P. is supported by the Fonds Erasme (Brussels, Belgium; Research Convention “Les Voies du Savoir”). L.T. was supported by the Fonds Erasme (Research Convention “Les Voies du Savoir”) and is supported by INNOVIRIS (Brussels, Belgium; research convention “DetectDem”). M.S. is Postdoctorate Clinical Master Specialist and X.D.T. is Clinical Researcher at the Fonds de la Recherche Scientifique (F.R.S.–FNRS; Brussels, Belgium). This research project is financially supported by INNOVIRIS (research convention “DetectDem”) and by a Stichting Alzheimer research grant. The MEG project at the HUB–Hôpital Erasme is financially supported by the Fonds Erasme (Research Convention “Les Voies du Savoir”) and the PET-MR project, by the Association Vinçotte Nuclear (AVN; Brussels, Belgium).

## Author contributions

V.P., X.D.T. and V.W. designed this study. L.T., F.A., S.D.W. acquired data. V.P. and V.W. developed analytical framework. V.P. performed coding and analyses. V.P., L.T., N.T., X.D.T. and V.W. analyzed and interpreted data. J.V.D., S.E., J-C.B., M.S. and X.D.T. provided resources. V.P. and V.W. wrote manuscript. All authors reviewed final version of manuscript.

## Supplementary Information

**Figure S1:**
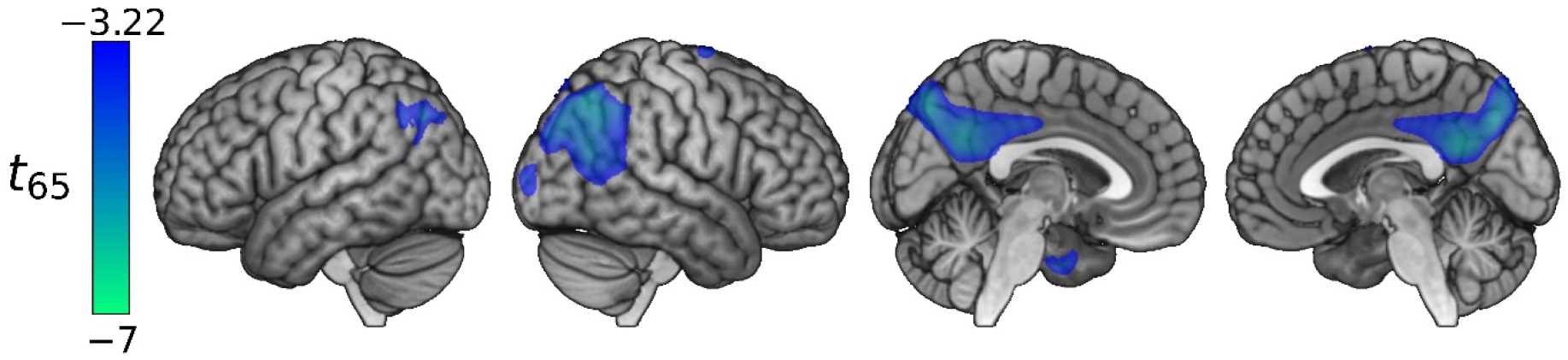
Cerebral glucose hypometabolism in AD patients. Negative statistical *t*_65_ contrast map of regional glucose metabolism in patients with AD (*n* = 43) vs. controls (*n* = 25), thresholded at *p <* 0.01 uncorrected (*t*_65_ *<* −3.22). The peaks at the left and right temporoparietal clusters and the precuneus cluster were also significant at *p <* 0.05, corrected for the family-wise error rate (threshold, *t*_65_ *<* −4.73). Age was included as covariate of non-interest. This analysis was performed using SPM12 (Wellcome Centre for Neuroimaging) on preprocessed FDG-PET images.

**Figure S2:**
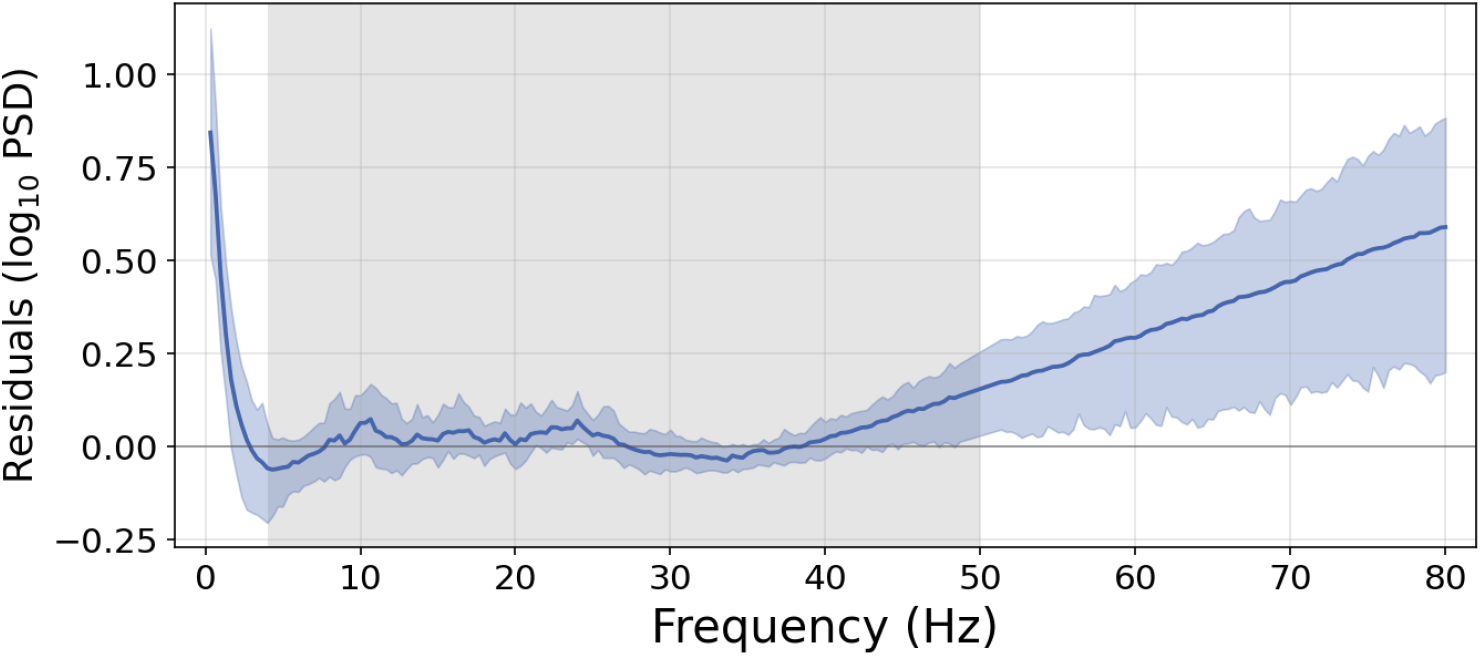
Residuals of the Lorentzian fit over 4–50 Hz. Residual spectrum (log_10_ PSD minus the fitted Lorentzian) as a function of frequency (group mean and 10th–90th percentile across participants). The Lorentzian was fitted in two steps over 4–50 Hz (shaded area), as in the main analysis, and oscillatory peaks (*α, β*) are masked. The flatness from 4 to about 40 Hz indicates adequacy of the Lorentzian fit. The sharp deviation below 4 Hz is due to the low-frequency power law (LRTC exponent), and the slow deviation from about 40 Hz, to a high-frequency noise power law. Power line bins (49–51 Hz) are not shown. Data are the mean MEG PSD over the bilateral primary sensorimotor cortices (precentral and paracentral Brainnetome parcels), computed without low-pass filtering, in patients with AD.

